# Dynamic Configuration of Coactive Micropatterns in the Default Mode Network during Wakefulness and Sleep

**DOI:** 10.1101/2020.07.29.226647

**Authors:** Yan Cui, Min Li, Bharat Biswal, Wei Jing, Changsong Zhou, Huixiao Liu, Daqing Guo, Yang Xia, Dezhong Yao

## Abstract

The default mode network (DMN) is a prominent intrinsic network that is observable in many mammalian brains. However, few studies have investigated the temporal dynamics of this network based on direct physiological recordings. Herein, we addressed this issue by characterizing the dynamics of local field potentials (LFPs) from the rat DMN during wakefulness and sleep with an exploratory analysis. We constructed a novel coactive micropattern (CAMP) algorithm to evaluate the configurations of rat DMN dynamics and further revealed the relationship between DMN dynamics with different wakefulness and alertness levels. From the gamma activity (40-80 Hz) in the DMN across wakefulness and sleep, three spatially stable CAMPs were detected: a common low-activity level micropattern (cDMN), an anterior high-activity level micropattern (aDMN) and a posterior high-activity level micropattern (pDMN). A dynamic balance across CAMPs emerged during wakefulness and was disrupted in sleep stages. In the slow-wave sleep (SWS) stage, cDMN became the primary activity pattern, whereas aDMN and pDMN were the major activity patterns in the rapid eye movement sleep (REM) stage. Additionally, further investigation revealed phasic relationships between CAMPs and the up-down states of the slow DMN activity in the SWS stage. Our study revealed that the dynamic configurations of CAMPs were highly associated with different stages of wakefulness and provided a potential three-state model to describe the DMN dynamics for wakefulness and alertness.

**Impact Statement:** In the current study, a novel coactive micropattern method (CAMP) was developed to elucidate fast DMN dynamics during wakefulness and sleep. Our findings demonstrated that the dynamic configurations of DMN activity are specific to different wakefulness stages and provided a three-state DMN CAMP model to depict wakefulness levels, thus revealing a potentially new neurophysiological representation of alertness levels. This work could elucidate the DMN dynamics underlying different stages of wakefulness and have important implications for the theoretical understanding of the neural mechanism of wakefulness and alertness.

## Introduction

Multimodal imaging studies of the human brain have discovered several intrinsic connectivity networks (ICNs) coexist during the resting state (Beckmann, DeLuca, Devlin, & Smith, 2005; Q. Liu, Farahibozorg, Porcaro, Wenderoth, & Mantini, 2017). Dynamic switching within these ICNs have demonstrated a hierarchical structure over time for brain activity at rest and have been significantly associated with cognitive traits (M. D. Fox et al., 2016; Vidaurre, Smith, & Woolrich, 2017). This suggests that brain activity is appropriately understood in terms of the dynamic configuration among ICNs. These studies have mainly considered each ICN as a whole during brain dynamics while ignoring the intrinsic dynamics of individual ICNs. Indeed, individual ICN also exhibits strong fluctuations in brain activity, and different ICNs are believed to dominate distinct cognitive functions (Rosazza & Minati, 2011). For a specific brain function, further tracking the dynamic configuration of fluctuations in brain activity at the single-ICN level might be critical for revealing the underlying physiological mechanism.

The default mode network (DMN) is one of the important ICNs, and is typically believed to be related to off-task internal mentations with high activity in the resting state (Gusnard, Akbudak, Shulman, & Raichle, 2001; M E Raichle et al., 2001). Recent studies have reported that the DMN is also engaged and displays positive contributions during several higher cognition task performances, such as the Tower of London task (D. Vatansever, Menon, Manktelow, Sahakian, & Stamatakis, 2015; Deniz Vatansever, Manktelow, Sahakian, Menon, & Stamatakis, 2018). Changes in wakefulness could also lead to alterations of DMN activity and connectivity in humans and rodents (K. C. R. Fox, Foster, Kucyi, Daitch, & Parvizi, 2018; Lu et al., 2012; Marcus E Raichle, 2015). DMN connectivity between the frontal and posterior areas in the human brain was reduced during the slow wave sleep (SWS) stage with a low level of wakefulness (Sämann et al., 2011). However, at sleep onset and throughout the rapid eye movement sleep (REM) stage, regions in the human DMN were shown to be persistently coupled (Horovitz et al., 2008; Larson-Prior et al., 2009). These findings illustrated that DMN activity was functionally reorganized during sleep and might further reflect levels of wakefulness and alertness. Additionally, fast and ever-changing dynamics of DMN activity have also been observed in various wakefulness levels in humans, implying that the temporal aspects of spontaneous DMN activity might be associated with alertness levels (Kapogiannis, Reiter, Willette, & Mattson, 2014; Panda et al., 2016). Thus, research investigating the dynamic configuration of DMN in different stages of wakefulness using direct physiological recordings is important.

The DMN could also be observable in rat brains, indicating its conservation in the mammalian brain during evolution (Huang et al., 2016; Lu et al., 2012). Though there exists several difference between the brain regions in rat DMN and human DMN, the anatomical topologies of them are visually similar (Hsu et al., 2016; Marcus E Raichle, 2015). In addition, neural activity of DMN regions also activated in awake rats, and suppressing the activity in the core DMN regions could modulate rats’ normal behavior (Tu, Ma, Ma, Dopfel, & Zhang, 2020; Upadhyay et al., 2011). Furthermore, the information flow within anterior and posterior DMN subsystems exhibited various alterations in different sleep stages and mental disorders in rats (Cui et al., 2018; Jing et al., 2017). The above findings imply that the DMN might carry a core function that transcends across species. Therefore, employing animal models to direct record physiological DMN signals is an effective way to investigate the temporal dynamics of DMN activity.

Several neurophysiological studies have reported that during the deep sleep stage, the neurons in various brain regions exhibited highly synchronized firing rates, occurring at approximately 0.5-2 Hz (Crunelli & Hughes, 2010; Gretenkord et al., 2020; Lőrincz et al., 2015). This specific activity pattern was identified as the up-down state, and has been considered a biomarker of low-level wakefulness in deep sleep (Jercog et al., 2017; Perez-Zabalza et al., 2020). Moreover, this up-down state has been demonstrated for both neuron membrane potentials and local field potentials (LFPs) (Holcman & Tsodyks, 2006) and characterizes the dynamics of slow oscillations during deep sleep (Ji & Wilson, 2007; Lőrincz et al., 2015). However, the existence of a physiological relationship between the up-down state and DMN dynamics is a topic of active interest, that deserves further exploration.

In the present study, we developed and applied a new dynamic activity pattern method to address these challenges. The new method, named the coactive micropattern analysis (CAMP), decomposed the fast dynamic activity into several intrinsic CAMPs and defined the brain dynamics through the constitutions and transitions among these CAMPs. We employed the CAMP analysis to elucidate the dynamic configurations of LFPs from rat DMN in different stages during wakefulness and sleep. Our results illustrate a reorganized dynamic configurations of CAMPs for fast DMN activity in different stages of wakefulness, implying that the dynamic configurations of DMN micropatterns might provide underlying neural correlates for the wakefulness levels observed during wakefulness and sleep.

## Methods and Materials

Detailed descriptions of the experimental procedures and data acquisition are described in the ***Supplementary Materials.*** Twenty-nine male Sprague-Dawley rats were used in our study. The DMN signals were acquired by chronically implanting fifteen electrodes into the brains of rats under deep anesthesia (Fig. 1 and Table 1). The rat DMN contained the following bilateral structures: the orbital frontal cortex (OFC), the rostral dorsal prelimbic cortex (PrL), the cingulate cortex (CG), the retrosplenial cortex (RSC), the dorsal hippocampus (HIP), the temporal lobe cortex (TE), the medial secondary visual cortex (V2) and the posterior parietal cortex (PPC). According to their anatomical coordinates (Lu et al., 2012), the PrL, OFC and CG regions were considered to be in the anterior subsystem of the DMN, whereas the RSC, HIP, PPC, TE and V2 regions were in the posterior subsystem of the DMN. In addition, we also implanted two electromyographic (EMG) electrodes bilaterally in the dorsal neck muscles. After DMN electrode implantation surgery, all rats recovered for approximately 2 weeks. During the recording session, the rats were placed in a noise-attenuated chamber and were allowed to move freely without anesthesia. All the signals for the LFP, the EMG and the videos signals were simultaneously recorded and continuously monitored for 72 h.

**Figure 1.**
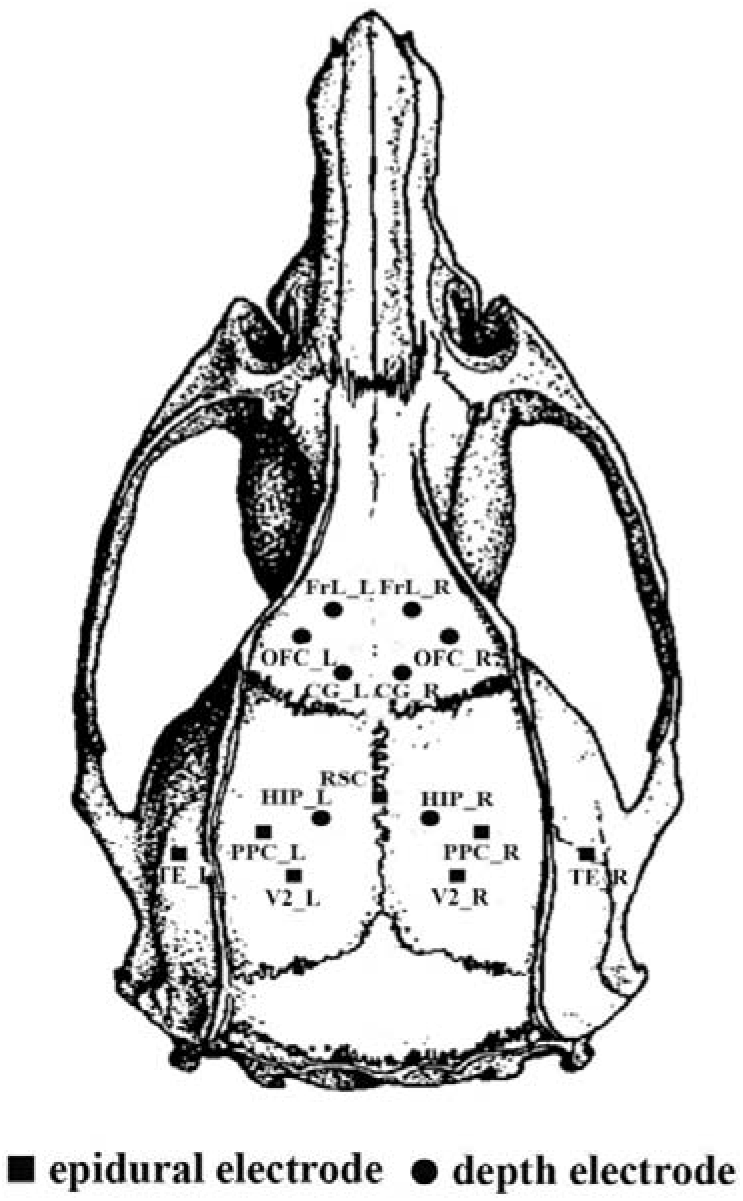
The placement of 15 intracranial electrodes.

The dataset used in the current study was selected from the last 24 h of the total recording and was separated into three stages, including resting (AWAKE), SWS and REM sleep stages. The AWAKE stage of rats was defined when the rats were standing or sitting quietly with low-amplitude and mixed-frequency LFP activity and relatively low and stable EMG activity. The SWS stage was the sleep duration when the rats were sleeping with high-amplitude and low-frequency LFP activity and low-level EMG activity. The REM sleep stage was the duration when the rats were sleeping with sawtooth-pattern LFP activity and flat EMG activity. For each rat, 30 segments in different stages were chosen, and each segment lasted 10 s (a total of 300 s of LFPs). All experimental animal procedures were approved by the Institutional Animal Care and Use Committee of the University of Electronic Science and Technology of China.

Moreover, we proposed a novel CAMP method to track fast DMN dynamics during wakefulness and sleep. Briefly, this method utilizes a point process approach that combines the advantages of both microstate analysis and coactive pattern analysis (X. Liu & Duyn, 2013; Michel & Koenig, 2018) and extracts CAMPs based on the extreme values of envelope signals at a high temporal resolution. Five steps were included in the CAMP algorithm. First, the original data were bandpass filtered into the gamma frequency band (40-80 Hz) and then Hilbert transformation was used to obtain the envelope signals (Fig. 2b). Second, the envelope signals were normalized and downsampled to improve the signal-to-noise ratio (SNR) for further analysis (Fig. 2c). Third, the active points for each envelope signal channel were then defined as the extreme points of the envelope signals, including local maximum and minimum values. Afterwards, the coactive patterns (CAPs) of the brain, which were defined as brain maps in which more than one brain region displayed active points at the same time point, were introduced for all stages (Fig. 2d). Fourth, a k-means clustering algorithm was applied to all the CAPs to decompose the CAMPs and the CAMP index (Fig. 2e). Finally, the criterion based on squared Euclidean distance was applied to update the CAMP and CAMP index (Fig. 2f). The last step was employed to precisely determine the final spatial structures of all CAMPs and the CAMP index. Using the CAMP method, we decomposed three stable CAMPs from gamma activity during DMN dynamics to reveal the fast changes in DMN activity in different stages of wakefulness. A detailed description of the CAMP analysis is provided in the ***Supplementary Methods***.

**Figure 2.**
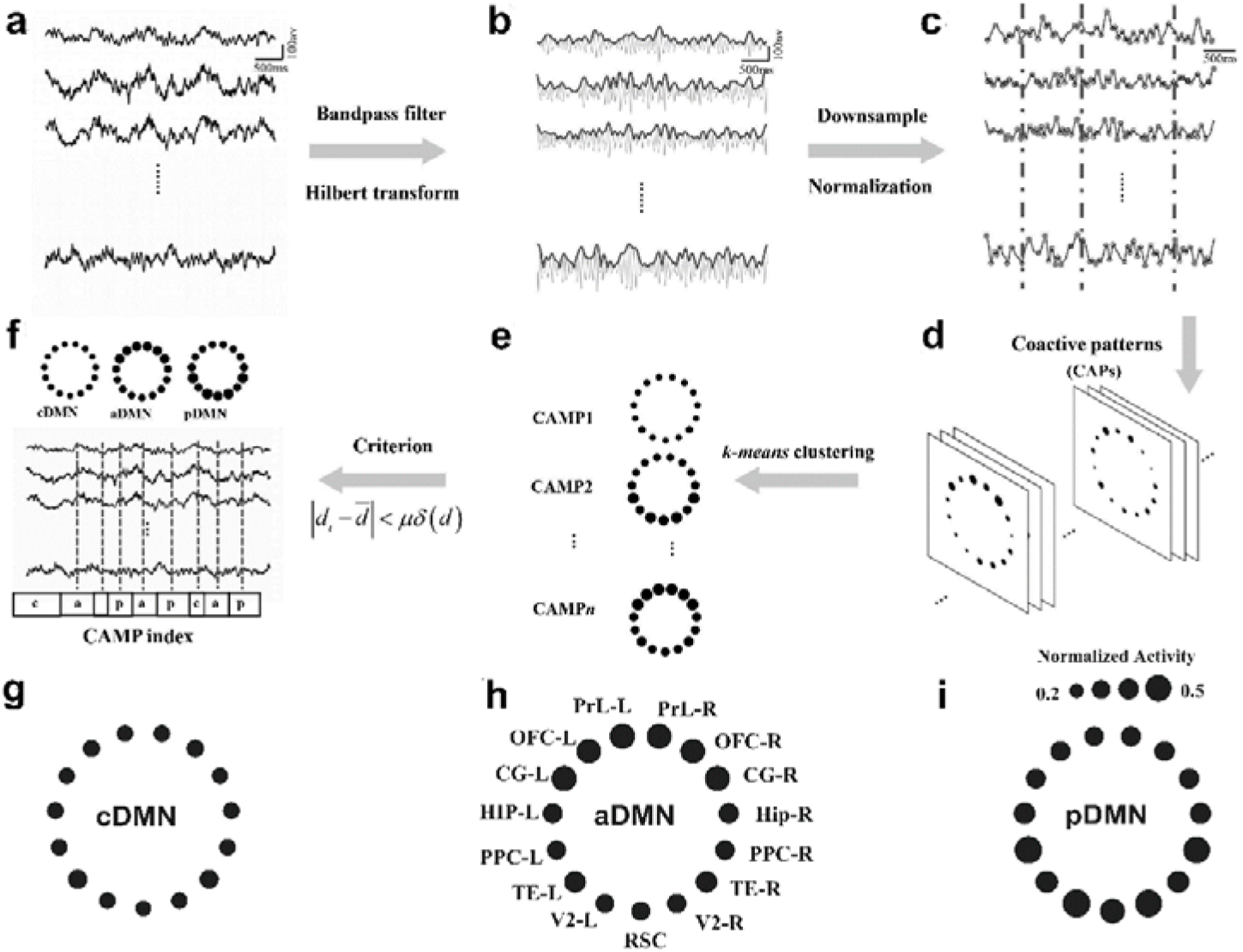
Schematic of the CAMP procedure and three CAMPs of gamma activity in the DMN during wakefulness and sleep. (a) The original LFPs. (b) The envelope signals (blue lines) were extracted by applying the Hilbert transform to the bandpass-filtered signals (gray lines). (c) All the envelope signals were downsampled (blue lines), and the extreme values were detected as the active points for each channel (red dots). The dotted lines suggest the coactive points in which more than N (N=7 in the present study) active points were observed across DMN regions. (d) The coactive patterns were the maps of activity of all DMN regions at coactive points. (e) The k-means clustering algorithm was applied to all coactive patterns to detect the CAMPs. (f) A criterion was employed to remove several coactive points and increase the aggregation of the CAMPs. The final CAMPs and CAMP index detected in this step were subjected to further analyses. (g) Spatial structure of the common low-activity level micropattern (cDMN). (h) Spatial structure of the anterior high-activity level micropattern (aDMN). (i) Spatial structure of the posterior high-activity level micropattern (pDMN).

The CAMP method was separately applied to the DMN activity of each segment in different stages for each rat and to the concatenated DMN activity of all rats and all stages during wakefulness and sleep. The CAMPs extracted from each rat in different stages were further employed to test their spatial stability across rats and wakefulness levels using Pearson correlation method. The results were derived from the CAMPs extracted from the concatenated DMN activities from all rats during wakefulness and sleep unless otherwise described.

## Results

### Three CAMPs of gamma activity in the DMN during wakefulness and sleep

The CAMP analysis procedure developed in the present study is schematically illustrated in Fig. 2 and described in more detail in the ***Supplementary Methods***. The concatenated gamma activities in the DMNs of all rats and all stages during wakefulness and sleep were decomposed into three distinct CAMPs, including a common low-activity level micropattern (cDMN), an anterior high-activity level micropattern (aDMN) and a posterior high-activity level micropattern (pDMN). In the cDMN, all DMN regions exhibited similar and low levels of activity (mean normalized activity: 0.2577 ± 0.0041, Fig. 2g), indicating a potential cooperation of them in this type of CAMP. However, two different levels of activity were observed in both the aDMN and pDMN with the aDMN exhibiting relatively higher levels of activity in the anterior DMN regions (mean normalized activity: 0.3868 ± 0.0018) and lower activity in the posterior DMN structures (mean normalized activity: 0.3050 ± 0.0060, Fig. 2h). In the pDMN, the posterior DMN structures displayed higher levels of activity (mean normalized activity: 0.3793 ± 0.0145), whereas the anterior DMN regions exhibited relatively lower levels of activity (mean normalized activity: 0.3073 ± 0.0021, Fig. 2i). Accordingly, both the aDMN and pDMN were considered high-activity micropatterns in DMN dynamics.

We separately decomposed the CAMPs for each rat in every wakefulness stage individually and tested their reliability across rats and stages. All three CAMPs exhibited high stability with large correlation coefficients among different rats during wakefulness and sleep (mean correlation coefficients: r = 0.7451, r = 0.7535, r = 0.6684 for the AWAKE, SWS and REM sleep stages, respectively; Table 2). In addition, the spatial structures of these CAMPs were also similar among the AWAKE, SWS and REM sleep stages (mean correlation coefficients: r = 0.6229, r = 0.7882, r = 0.8600 for the cDMN, aDMN and pDMN, respectively; Table 3). These findings demonstrated the high reliability and robustness of these CAMPs.

### Temporal features and activity levels of each CAMP during wakefulness and sleep

We computed several temporal measurements, including the total occurrence (occurrence probability), total duration (duration probability) and mean duration, to characterize the features and dynamics of these CAMPs during wakefulness and sleep. All these features represented the temporal properties of these CAMPs in different stages. Based on the comparisons, all features of cDMN displayed the largest values in the SWS stage and the smallest values in the REM sleep stage, and the two high-activity micropatterns (aDMN and pDMN) exhibited the largest values for all features in the REM sleep stage and the smallest values in the SWS stage (Fig. 3a-3c). These opposite alterations in features between low- and high-activity micropatterns suggests that these two types of CAMPs might play different physiological roles for wakefulness. Besides, all the features in three stages were remarkably different among CAMPs, improving our knowledge of the changes in wakefulness levels during wakefulness and sleep.

**Figure 3.**
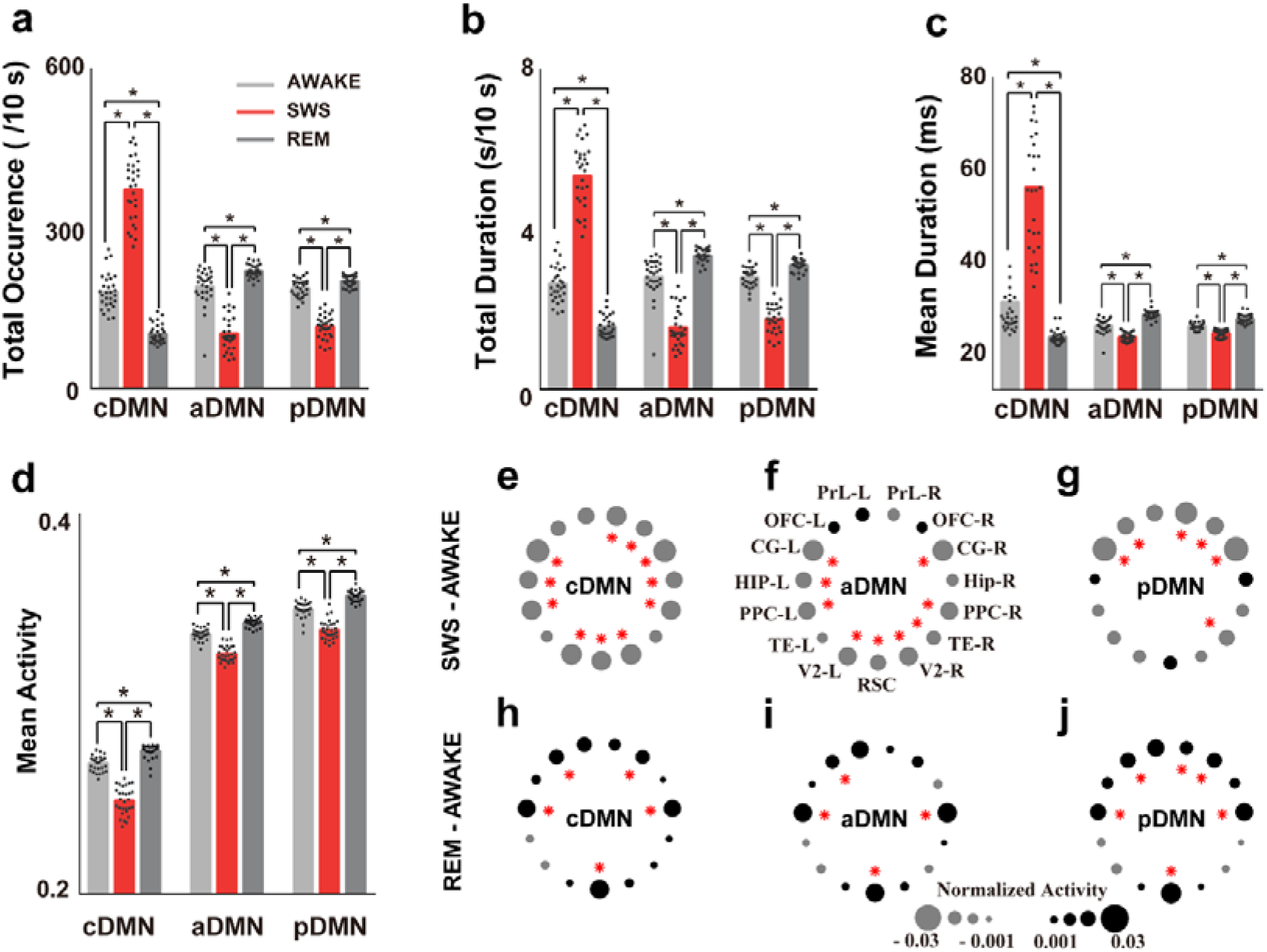
Comparisons of the temporal features and activity levels of each CAMP during wakefulness and sleep. (a) Comparisons of the total occurrence of each CAMP in different stages of wakefulness. The dots represent the values obtained from 29 rats, and the black stars indicate significant differences with a corrected p<0.001. (b) Comparisons of the total duration. (c) Comparisons of the mean duration. (d) Comparisons of the mean DMN activity during wakefulness and sleep for different CAMPs. (e-j) Comparisons of activity in DMN nodes for different CAMPs across different stages of wakefulness: (e and h) cDMN, (f and i) aDMN, and (g and j) pDMN. Gray dots indicate decreased normalized activity and black dots indicate increased normalized activity. The size of the dot reflects the value of the difference, and the red stars indicate significance differences with a corrected p<0.001.

However, all of these CAMPs displayed different activities in DMN regions during wakefulness and sleep. In particular, all DMN regions exhibited reduced activity during SWS stages in all CAMPs (Fig. 3e-3g). Moreover, the posterior DMN structures exhibited significantly reduced activity in the aDMN, while the anterior DMN regions exhibited significantly reduced activity in the pDMN. All of these regions showed relatively lower activity in the AWAKE stage, indicating a preservation of the major activity in these two micropatterns during deep sleep (Fig. 3f-3g, red stars). However, all CAMPs displayed increased activity in most DMN regions during the REM sleep stage. The activities in the HIP, OFC and RSC regions were significantly increased during the REM sleep stage in all CAMPs, implying the importance of these DMN regions for REM sleep (Fig. 3h-3j, red stars). In addition, the mean activity level of each CAMP exhibited similar variation trends across different stages of wakefulness. The lowest mean activity of CAMPs was observed in the SWS stage, whereas the highest mean activity was observed in the REM sleep stage (Fig. 3d).

### The features and transitions of CAMPs during wakefulness and sleep

The configurations of these CAMPs involved in DMN dynamics in different stages were also distinct (Fig. 4a-4c). All CAMPs presented similar features in the AWAKE stage (occurrence probabilities: 32.12%, 34.12% and 33.75%; duration probabilities: 31.68%, 34.15% and 34.17%; and mean duration: 29.79 ms, 24.41 ms and 24.37 ms for the cDMN, aDMN and pDMN, respectively). No significant differences of the features among three CAMPs were observed, indicating that their roles were equivalent and that a dynamic balance in DMN activity might exist among CAMPs at wakeful rest. However, the cDMN became the dominant activity pattern of DMN dynamics in the SWS stage as it had the largest occurrence probabilities (62.66%, 17.56% and 19.78% for the cDMN, aDMN and pDMN, respectively), duration probabilities (61.47%, 18.02% and 20.51% for the cDMN, aDMN and pDMN, respectively) and mean duration (55.58 ms, 21.89 ms and 22.78 ms for the cDMN, aDMN and pDMN, respectively) among all CAMPs. The predominant constituent of the low-activity micropattern suggests that all the DMN regions might have been in a stage of low activity and that DMN activity preferred a silent pattern during deep sleep. However, the two high-activity micropatterns were the main CAMPs during the REM sleep stage. All the features of aDMN and pDMN were significantly larger than those of cDMN (occurrence probabilities: 19.42%, 42.05% and 38.53%; duration probabilities: 19.33%, 41.84% and 38.83%; and mean duration: 21.88 ms, 26.92 ms and 26.01 ms for the cDMN, aDMN and pDMN, respectively). The greater percentage of high-activity micropatterns during REM sleep suggests a reactivation of DMN activity in this stage. In addition, comparisons of features within two high-activity micropatterns demonstrated that the aDMN displayed significantly larger values for the three features, implying that it played a more important role in REM sleep.

**Figure 4.**
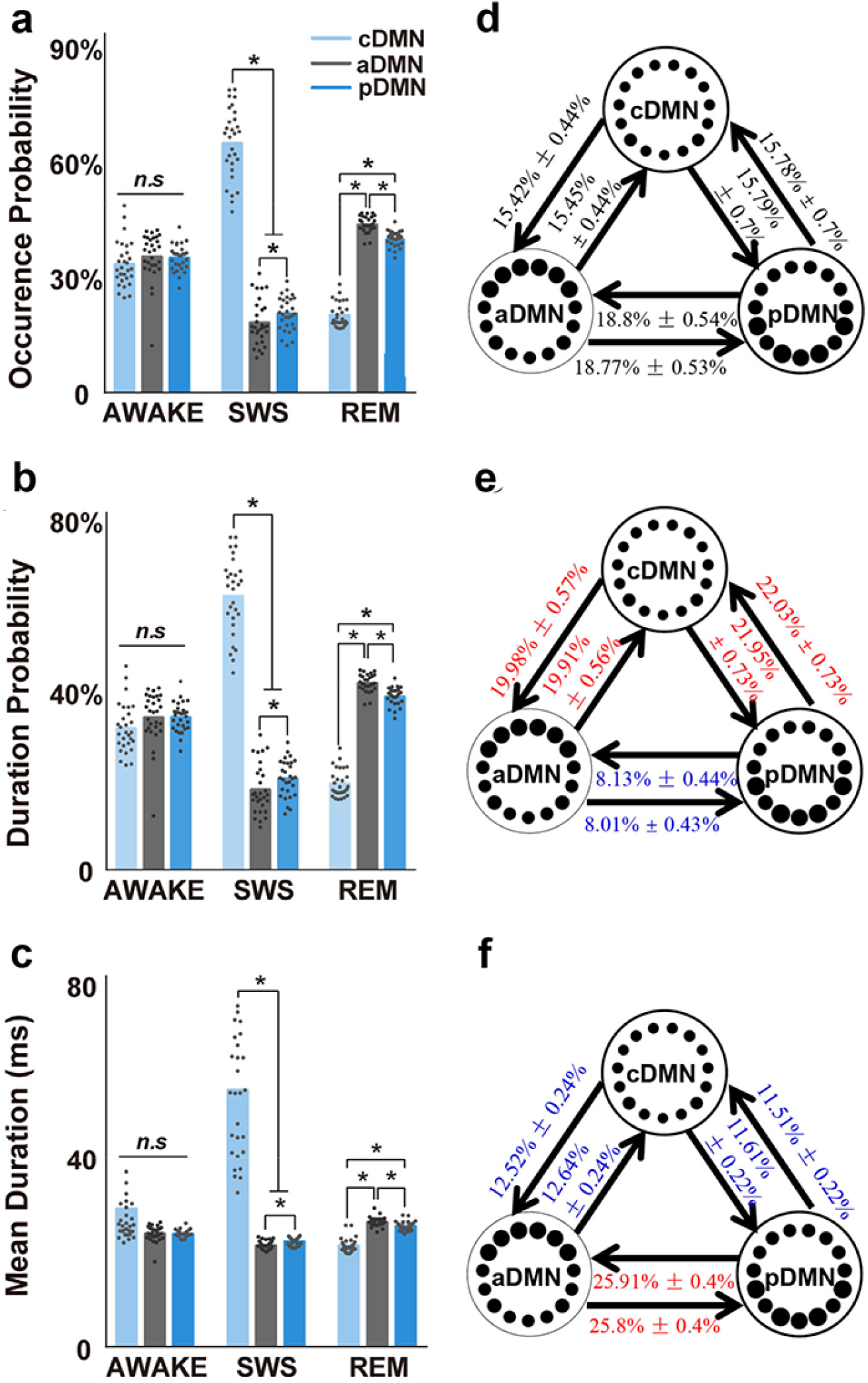
Characteristics of CAMPs and the transitions among them in different stages of wakefulness during wakefulness and sleep. (a) Comparisons of the occurrence probability for all CAMPs in the three stages. The black dots indicate the values of the occurrence probability obtained from 29 rats in different CAMPs and stages. The black stars indicate significant differences with a corrected p<0.001. (b) Comparisons of the duration probability. (c) Comparisons of the mean duration. (d-f) The transition structures among CAMPs for the AWAKE (d), SWS (e) and REM sleep stages (f). All the numbers indicate the mean TPs calculated for the 29 rats and the standard deviation. The numbers in blue indicate a significantly lower transition probability than observed in the AWAKE stage, and the numbers in red indicate a significantly higher transition probability. The significance level is a corrected p<0.001.

Furthermore, the temporal concatenations of these CAMPs (i.e., the CAMP indices) in different stages also exhibited specific changes. We first performed a randomization test to examine the transition structures of these CAMP indices in different stages. The transitions among CAMPs occurred randomly in the AWAKE stage (p = 0.8157), indicating that the transition probabilities (TPs) of pairs of CAMPs in the resting stage were proportional to their occurrences. However, these transitions did not occur randomly in the SWS (p<0.0001) or REM sleep stages (p<0.0001), suggesting the stabilization of the CAMP index structures during the sleep cycle and further implying the existence of several preferred transitions among CAMPs in both SWS and REM sleep stages.

Next, we compared the TPs for pairs of CAMPs between the two sleep stages and the AWAKE stage. We revealed similar TPs in the AWAKE stage (no significant differences among all TPs, Fig. 4d), suggesting the presence of balanced transitions among all CAMPs at rest. However, the TPs within the two high-activity micropatterns showed significant reductions in the SWS stage, whereas those between the high-activity micropatterns and the low-activity micropattern increased significantly (Fig. 4e). These changes in TPs emphasized the functional role of inhibitory activity in DMN regions in deep sleep. On the other hand, TPs in the REM sleep stage displayed different alterations, including significantly increased TPs within the high-activity micropatterns and remarkable decrease in TPs between the high-activity micropatterns and the low-activity micropattern (Fig. 4f). The increased transitions within two high-activity micropatterns revealed activation of DMN regions during REM sleep. Based on these findings, the CAMP indices and the functional roles of these CAMPs were specific for different stages. The alterations in DMN activity during wakefulness and sleep might be attributed to the specific temporal combinations of the CAMPs constituting the activity in different stages rather than the spatial structures of CAMPs themselves, which were rather stable across different stages.

### Strong phasic relationships between CAMPs and up-down states in the SWS stage

Up-down states are considered the predominant pattern of slow oscillations (0.5-2 Hz) during the SWS stage. Estimations of the phase distributions of each CAMP in the anterior and posterior DMN slow activity regions with the Hilbert transformation demonstrated that these CAMPs displayed strong phasic relationships with the up-down states in the SWS stage. The cDMN preferred the down state of anterior DMN activity (Fig. 5a, significant directionality: 1.97 π, red line) and the up state of posterior DMN activity (Fig. 5d, significant directionality: 1.16 π, red line). Additionally, both the aDMN and pDMN were phase locked to the up state of anterior DMN activity (Fig. 5b, significant directionality: 1.21 π for aDMN; Fig. 5c, significant directionality: 1.18 π for pDMN) and the down state of posterior DMN activity (Fig. 5e, significant directionality: 0.23 π for aDMN; Fig. 5f, significant directionality: 0.18 π for pDMN). These similar phasic relationships implied that two high-activity micropatterns might belong to the same activity pattern of slow oscillations during deep sleep. Thus, our proposed CAMPs may reflect the up-down states of DMN slow activity in the SWS stage, and a close physiological association existed between the up-down states with DMN dynamics.

**Figure 5.**
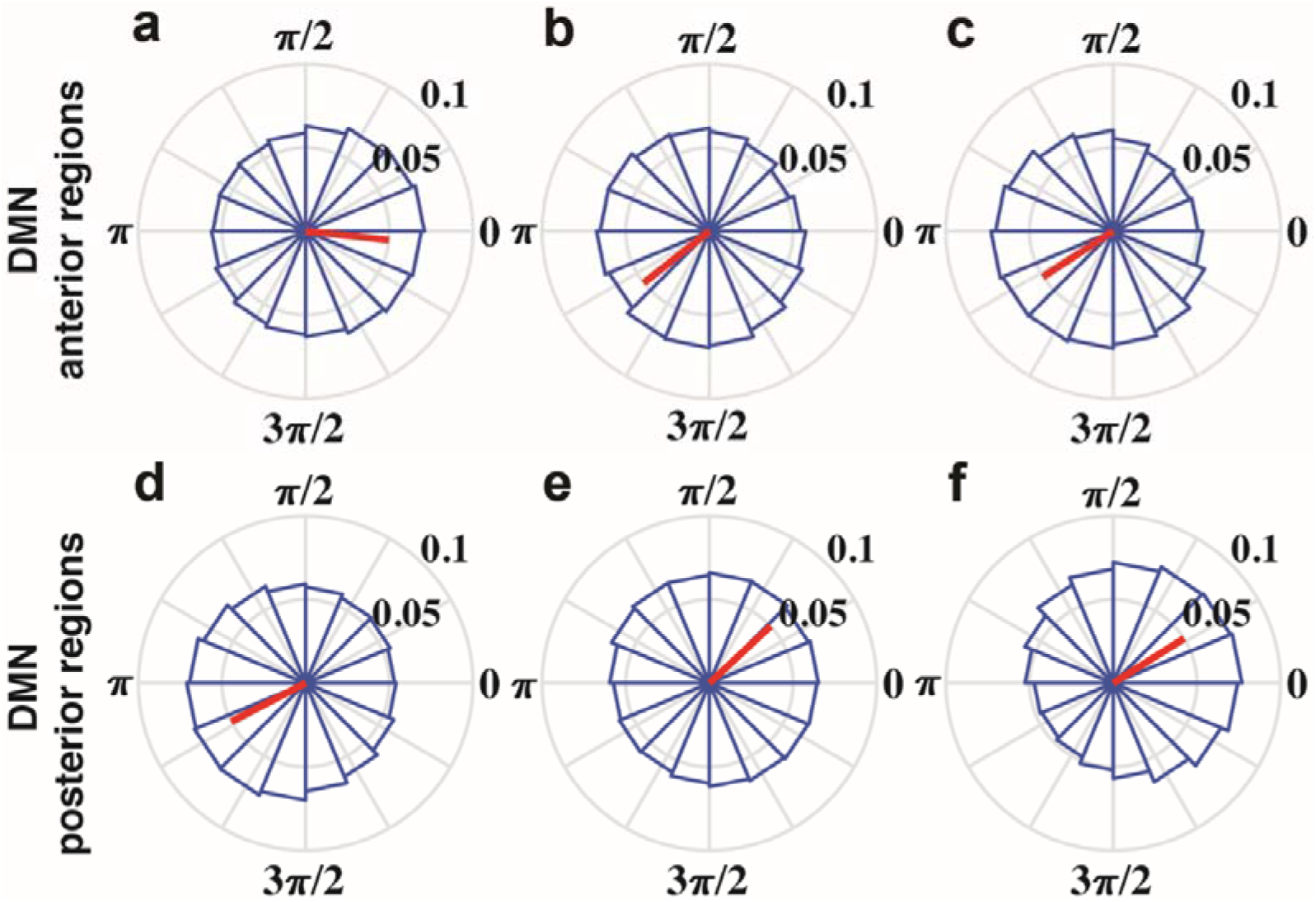
Phase locking relationship between each CAMP with slow oscillations in the SWS stage. (a-c) The phase locking relationships between the cDMN (a), aDMN (b) and pDMN (c) with the slow oscillations in anterior DMN regions. (d-f) The phase locking relationships between the cDMN (d), aDMN (e) and pDMN (f) with the slow oscillations in posterior DMN regions. The red lines showed the significant directionality with Rayleigh test p<0.001.

## Discussion

In the present study, we proposed a coactive micropattern (CAMP) algorithm to reveal the dynamics of DMN based on the direct physiological recordings in rat during wakefulness and sleep. Our results indicated that the fast dynamics of DMN gamma activity could be decomposed into three different CAMPs. These CAMPs exhibited stable spatial structures across wakefulness and sleep, while their dynamic configurations were specific to different stages. In addition, all these CAMPs were strongly phase locked to the up-down states of slow DMN activity in the SWS stage, suggesting the temporal sequence of the neural relationship between up-down states and DMN dynamics. Our findings described the distinct dynamic configurations of DMN activity during wakefulness and sleep. The proposed a three-state model may reveal a neural mechanism by which DMN dynamics mediated wakefulness and alertness.

### Physiological significance of the three CAMPs

Previous studies have reported a strong correlation between electrophysiological gamma activity and blood oxygen level-dependent (BOLD) signals (N. K. Logothetis, Pauls, Augath, Trinath, & Oeltermann, 2001; Nikos K. Logothetis, 2002; Magri, Schridde, Murayama, Panzeri, & Logothetis, 2012; Scheering, Koopmans, Van Mourik, Jensen, & Norris, 2016). In addition, DMN regions have also shown deactivation at gamma frequency during the performance of external tasks in several human electroencephalography (EEG) studies (Karim Jerbi et al., 2010; Ossandon et al., 2011), indicating the importance of gamma oscillation in DMN activity. Hence, we specifically focused on the fast dynamics of DMN gamma activity in the current study. The gamma activity in the rat DMN was decomposed into three stable CAMPs during wakefulness and sleep. We also showed the CAMPs decomposed from DMN alpha (8-13Hz) and beta (13-30Hz) activity, which exhibited similar structures with those found in gamma band (Supplementary Fig. 5). The differences across these CAMPs further provided direct electrophysiological evidence that the DMN regions might not be simultaneously activated. Besides, different CAMPs had distinct mean durations. These phenomena revealed the differences in the activation times of anterior and posterior DMN structures in the fast dynamics and further illustrated the diversity in the latencies for both the excitation and inhibition of DMN regions (Brett L. Foster, Mohammad Dastjerdi, 2012; Foster, Rangarajan, Shirer, & Parvizi, 2015).

Indeed, both human and animal studies found that the DMN structure could be separated into two subnetworks, i.e., a parietal subnetwork and a prefrontal subnetwork (Cui et al., 2018; Hagmann et al., 2008; Lu et al., 2012; Wu et al., 2017). In the present study, we not only reinforced this finding from the aspect of fast DMN dynamics but also provided a possible dynamic substrate for this separation of the DMN structure. As a key component of the DMN, the orbital frontal cortex (OFC) has historically been posited to integrate interoceptive and exteroceptive information from multisensory stimuli to process information about the internal and external bodily milieu (Ongur & Price, 2000). Accordingly, we hypothesized that the high-activity micropattern aDMN might play an important role in making inferences and guiding actions in a timely and environmentally relevant manner.

Furthermore, the retrosplenial cortex (RSC), another key area in the DMN, has extensive connections with the hippocampal formation. The projections between the RSC and hippocampal formation provide an important pathway that regulates learning, memory and emotional behavior (Wyss & Vangroen, 1992). Furthermore, the hippocampal formation is a limbic structure that forms direct or indirect connections to other DMN regions. Therefore, the high-activity micropattern pDMN detected in the present study might be associated with memory and emotional behavior. Additionally, both the aDMN and pDMN were strongly phase locked to the up state of anterior DMN activity and the down state of posterior DMN activity during the SWS stage, indicating that they may reflect similar performances for the up-down states of slow oscillations during DMN dynamics. Moreover, these two high-activity micropatterns together accounted for more than 70% of the time in the resting state, which helps explain why the brain requires high basal cerebral blood flow and metabolism for spontaneous activity (Marcus E Raichle & Mintun, 2006). It should be noted that the DMN structure could also be split into dorsal and ventral branches according to the dorsal and medial temporal regions of RSC and hippocampus in human brain (Chen, Glover, Greicius, & Chang, 2017; Shirer, Ryali, Rykhlevskaia, Menon, & Greicius, 2012). However, the rat DMN is commonly divided into the anterior and posterior subsystems for the anatomical difference with human DMN (Marcus E Raichle, 2015).

We also observed a low-activity micropattern (i.e., cDMN) in DMN dynamics that was widely distributed in all wakefulness stages. In the cDMN, all DMN regions displayed lower activity, indicating that the cDMN could represent the silent state for DMN activity in which all the DMN regions prefer relaxations and are prepared for the next excitation. Moreover, the cDMN was the only coactive micropattern in which all DMN regions operated in the same manner in DMN dynamics. Thus, the appearance of cDMN suggested a working mode for DMN with low energy, but this concept requires further study.

### The balance of dynamic DMN configurations supports wakefulness and alertness during wakefulness

Based on accumulating evidence, DMN activity is tightly correlated with wakefulness levels in health and disease (Buckner, Andrews-Hanna, & Schacter, 2008; Kapogiannis et al., 2014; Panda et al., 2016). In the AWAKE stage, all the CAMPs exhibited similar features, and the dynamic transitions among them were not significantly different. These similarities illustrated a balanced dynamic configuration among these CAMPs during fast gamma activity in the DMN at rest. The DMN is a key network involved in integrating high-order information from multiple sensory modalities based on numerous projections from variable somatic cortex and core limbic structures (HIP and amygdala) to the DMN regions (Heidbreder & Groenewegen, 2003; Reep, Chandler, King, & Corwin, 1994). These projections might provide the anatomical substrate for the correlation of DMN activity with alertness levels, which are largely believed to be determined by global levels of arousal regulated by the brainstem via the reticular activating system (RAS) (Delano-Wood et al., 2015). Accordingly, the identified balance of DMN dynamics might be a competitive product between the integration and differentiation of DMN activity in maintaining wakefulness and alertness during resting state (Cavanna, Vilas, Palmucci, & Tagliazucchi, 2018; Tononi, 2004). Furthermore, this balance of dynamic configurations also indicated that the DMN might function in multistable regimes and revealed the potential neural mechanism by which DMN activity supported wakefulness and alertness in the resting state (Andrews-Hanna, 2012; Buckner et al., 2008).

### Functional reorganization of dynamic DMN configurations during sleep

Compared to the resting state, the SWS stage was consistently accompanied with reduced brain activity, whereas commensurate brain activity has been reported in the REM sleep stage (Horovitz et al., 2008). Consistent alterations in the average brain activity associated with CAMPs during DMN dynamics were also observed in our study, suggesting that the activities of CAMPs might also reveal the changes in wakefulness during wakefulness and sleep. However, the reduced activities of all CAMPs might not sufficiently explain the decrease in DMN activity observed during deep sleep due to the stable spatial structures of these CAMPs during wakefulness and sleep. The decrease in activity might result from the increased occurrence probability of the cDMN and the decreased probabilities of other two high-activity CAMPs. These inversely changed occurrence probabilities in different CAMPs revealed the neural mechanism of reduced brain activity given that the DMN regions were shown to prefer the low-activity state during deep sleep (Bazhenov, Timofeev, Steriade, & Sejnowski, 2002; Diekelmann & Born, 2010).

The balance of dynamic DMN configurations was also disrupted during sleep, indicating the functional reorganization of DMN dynamics. The reorganization of DMN activity might be associated with the alteration of different wakefulness levels in different sleep stages (Tononi, 2004). In the REM sleep stage, the dynamic transitions between the aDMN and pDMN increased, indicating more communication between anterior and posterior DMN regions. The communications displayed the top-down and bottom-up mechanisms in DMN structure, both of which are important for the information processing in the brain (Buschman & Miller, 2007; Theeuwes, 2010). Thus, we speculate that the communications between anterior and posterior DMN regions might help elucidate the neurophysiological basis underlying the preservation of the wakefulness level in the REM sleep stage.

In the SWS stage, the dynamic transitions between the low-activity micropattern and two high-activity micropatterns increased significantly. Moreover, different types of micropatterns corresponded to distinct up-down states in slow oscillations among DMN regions. Accordingly, the transitions between the low-activity micropattern and two high-activity micropatterns in DMN dynamics could be deemed as the transitions within up-down states. The dominant transitions of up-down states in deep sleep further suggested the physiological importance of these increased dynamic transitions. However, the dynamic transitions within the two high-activity micropatterns decreased in the SWS stage. These reductions supported our hypothesis that communications between anterior and posterior DMN regions are important for levels of wakefulness and alertness given that wakefulness and alertness are almost lost during deep sleep. The loss of wakefulness and alertness might not be caused by the change in a single type of dynamic transition within pairs of CAMPs. We hypothesized that the balance of dynamic DMN configurations was the underlying key neural mechanism supporting wakefulness and alertness, which emerged during wakefulness and disappeared during sleep. The coordination and cooperation of all CAMPs played a core role for the ability of the DMN in supporting wakefulness and alertness.

Based on these findings, we propose a three-state model to describe the relationship between DMN micropatterns and wakefulness levels observed during wakefulness and sleep. As shown in Fig. 6, the three CAMPs involved in DMN dynamics are the basis of this model, and their interactions refer to the underlying mechanism regulating the wakefulness level observed in distinct stages. Equal communications among the three CAMPs support conscious awareness in the AWAKE stage. The communications between the low-activity micropattern (i.e., cDMN) and each high-activity micropattern (i.e., aDMN and pDMN) are important for the SWS stage characterized by a low level of wakefulness. During the REM sleep stage, communications within high-activity micropatterns are predominant.

**Figure 6.**
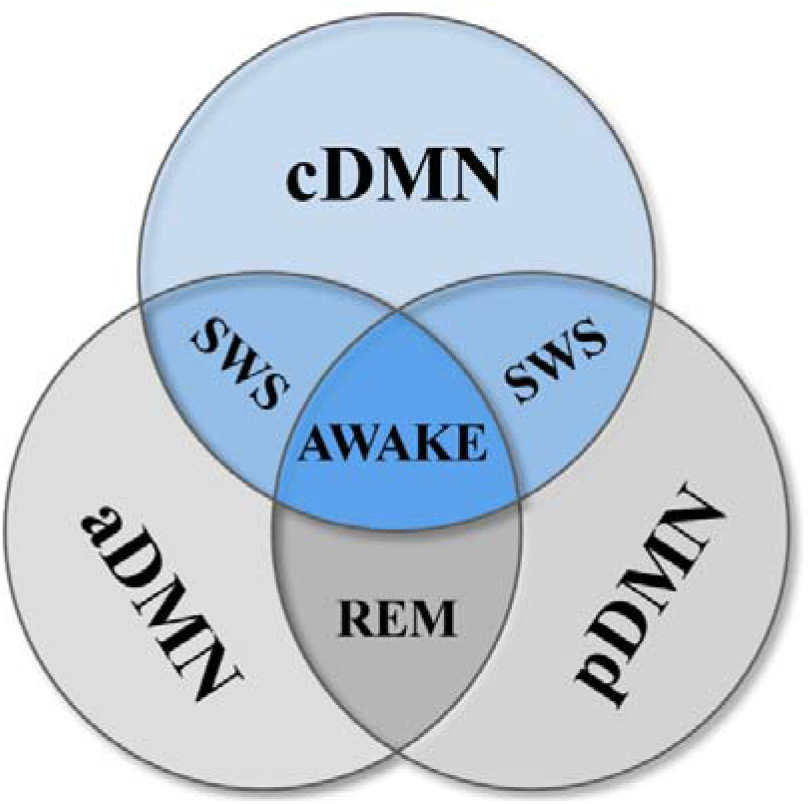
The three-state model of the alertness levels during wakefulness and sleep. The AWAKE stage requires the cooperation of all three CAMPs, while the SWS stage requires communications between the low-activity micropattern (cDMN) and the high-activity micropatterns (aDMN or pDMN). The REM sleep stage requires interactions within the two high-activity micropatterns.

According to the proposed three-state model, we hypothesize that preservation of wakefulness and alertness not only requires information transition between anterior and posterior DMN regions, but also need a state that all DMN regions remain silent and relaxed. Information transition within anterior and posterior DMN regions is mediated by up-down and bottom-up mechanisms and vital for supporting wakefulness and alertness. The absence of this process could lead to the loss of alertness in the SWS stage, and this process alone would result in the wakefulness level of the REM sleep stage, which is more wakeful than the SWS stage and less wakeful than the AWAKE stage. This phenomenon highlights the importance of the silent pattern for all DMN regions during the resting state with wakefulness and alertness.

Wakefulness and alertness in humans not only depend on the anterior-posterior integration with DMN regions but also involves fronto-parietal task-positive executive and attention networks. Moreover, the connections between DMN and task positive networks also play critical roles in supporting wakefulness and alertness. Given the limitation of neuroimaging measure, the current work only describes the associations between DMN dynamics and wakefulness levels. The roles of integrations between DMN and other networks could not be explored at present. Further work could validate our model with DMN activity during the wakefulness-sleep cycle and further track the roles of integrations among different ICNs across different levels of wakefulness and alertness with human EEG signals.

### Methodological perspectives

Consistent with the promising microstate analysis of EEG/LFP signals (Michel & Koenig, 2018), the CAMP analysis reported herein also assumes that brain activity consists of several distinct instantaneous patterns. The difference is that the CAMP method focuses on the nature of brain activity in different regions and extracts micropatterns from the envelope signals. Envelope signals imply temporal alterations of brain power, and their decomposition directly reveals brain rhythm dynamics. In addition, the coactive patterns analyzed in the CAMP method were selected based on the distributions of extreme values in the envelope signals, which differs from the method used in microstate analysis. Local extreme values in envelope signals represent the instantaneous higher/lower activities of brain regions followed by contrasting changes in activity. The derived coactive patterns were thus considered to represent the activity patterns leading to inversion of activity among regions in specific brain networks. Therefore, we postulate that the proposed CAMP method will help researchers elucidate coactive micropatterns in specific brain networks and reveal additional underlying information about fast brain dynamics.

### Limitations

Although we reveal several interesting findings in the current work, some limitations exist that should be taken into consideration. First, the rat DMN is slightly anatomically different from the human DMN, and our findings in rat DMN dynamics need to be validated in the human DMN during wakefulness and sleep. In addition, the present work reveals the dynamic configurations of DMN activity exclusively in the gamma band, and different frequency bands have distinct physiological roles. The relationships between the DMN dynamics in other frequency bands with wakefulness levels should be investigated in future studies.

## Conclusion

The DMN is believed to be associated with neural mechanisms of wakefulness and alertness levels, whereas fast dynamics of DMN activity based on direct physiological recordings in different stages of wakefulness remain unclear. We highlighted that the fast dynamics of DMN activity during wakefulness and sleep shared structurally stable CAMPs, whereas their dynamic configurations were specific to different levels of wakefulness. Our results indicated the reorganization of DMN dynamics during wakefulness and sleep, and provided a three-state model to reveal the fundamental neural associations between DMN activity and wakefulness levels.

## Acknowledgments

We thank Professor Pedro A. Valdes-Sosa for the valuable discussions and suggestions regarding this work.

## Authorship Confirmation Statement

Y.X., D.Y., C.Z., and D.G. designed the research; Y.C., M.L., and W.J. performed the research; Y.C., M.L., B.B., and H.L. analyzed the data; and C.Y., B.B., D.G., and D.Y. wrote the paper.

