## Supplementary material for "Dynamic Configuration of Coactive Micropatterns in the Default Mode Network during Wakefulness and Sleep": Tables

|  | **A-P** | **M-L** | **D-V** |
| --- | --- | --- | --- |
| **PrL** | 4.2 | ± 0.8 | 3 |
| **OFC** | 3.7 | ± 1.8 | 4.7 |
| **CG** | 1.7 | ± 0.7 | 2.6 |
| **RSC** | -3.3 | 0 | 0 |
| **HIP** | -4.3 | ± 1.4 | 3 |
| **PPC** | -4.5 | ± 4 | 0 |
| **V2** | -5.2 | ± 2.4 | 0 |
| **TE** | -5.2 | ± 8 | 5 |

**Table 1.** Coordinates of the 15 electrodes (mm). A-P, M-L, and D-V indicate anterior-posterior, medial-lateral, and dorsal-ventral directions, respectively**.** PrL, prelimbic cortex; OFC, orbital cortex; CG, cingulate cortex; RSC, retrosplenial cortex; HIP, hippocampus; PPC, posterior parietal cortex; V2, secondary visual cortex; TE, temporal association cortex.

|  | **cDMN** | **aDMN** | **pDMN** | **Mean Reliability** |
| --- | --- | --- | --- | --- |
| **AWAKE** | 0.5806 | 0.8043 | 0.8504 | 0.7451 |
| **SWS** | 0.7588 | 0.6472 | 0.8544 | 0.7535 |
| **REM** | 0.4257 | 0.6951 | 0.8843 | 0.6684 |

**Table 2.** The reliabilities (correlation coefficients) of CAMPs in the three stages across rats.

|  | **AWAKE** *vs.* **SWS** | **AWAKE** *vs.* **REM** | **SWS** *vs.* **REM** | **Mean Reliability** |
| --- | --- | --- | --- | --- |
| **cDMN** | 0.7214 | 0.6352 | 0.5122 | 0.6229 |
| **aDMN** | 0.7910 | 0.8650 | 0.7087 | 0.7882 |
| **pDMN** | 0.8406 | 0.9091 | 0.8302 | 0.8600 |

**Table 3.** The reliabilities (correlation coefficients) of CAMPs across the three conscious stages.
