## Supplemental Information for "Dynamic Configuration of Coactive Micropatterns in the Default Mode Network during Wakefulness and Sleep"

**Supplementary Information**

SI Materials

**Dataset.** Twenty-nine male Sprague-Dawley rats were used in our experiment. Firstly, fifteen electrodes, including seven epidural cortical electrodes and eight depth electrodes, were implanted into the brain of each rat under deep anesthesia (sodium pentobarbital, 60 mg/kg body weight, i.p.) at the coordinates proposed by Lu(Lu et al., 2012) (Fig. 1, Table 1). The reference electrode was placed in the cerebellum, and two electromyographic (EMG) electrodes were implanted bilaterally in the dorsal neck muscles. Here the cerebellum was chosen for the placement of the reference electrode, for that there was lower neural activity in the cerebellum and the cerebellum was not involved in many cognitive functions. Notably, 0.6 ml of atropine sulfate (0.5 mg/ml, s.c.) was injected during electrode implantation to prevent excess secretions from the respiratory tract. Meanwhile, the body temperature of the rats was maintained at 37 degrees centigrade with a heating pad. Then, all electrodes were welded to connectors and fixed on the skull of the rat with dental acrylic. After the surgical procedure, penicillin G was administered to prevent infection, and all rats were allowed at least 2 weeks for recovery before the recording session started. All experimental animal procedures were approved by the Institutional Animal Care and Use Committee of the University of Electronic Science and Technology of China.

Prior to the recording sessions, the rats were habituated to the experimental environment and the recording cable for 2 days. During the recording session, all rats were placed in a glass box on a 12-h light/dark cycle (lights on at 8:00 am). Each recording electrode was connected to an acquisition system (Chengyi, RM62160, China). Electrophysiological signals (LFPs) and videos were synchronously and continuously acquired for 72 h. The amplified and filtered (0.16–100 Hz for LFPs, 8.3–500 Hz for electromyogram (EMG), and 50-Hz notch filter) signals were stored on a hard disk (Lenovo Company, USA), and the sample frequency was set to 1,000 Hz. All experiments were performed in a noise-attenuated room, where the background noise was set to 32.2 ± 3.0 dB and the temperature was maintained at 25 ± 0.5 degrees centigrade. The experimenter entered the noise-attenuated room to replace food and water and clean cages at 12:00 am daily.

After the end of experiments, the histological images of DMN regions for each rat were performed to check whether the positions of electrodes were right in those DMN regions, especially for eight depth regions. The histological images for PrL, OFC, CG and HIP were showed in Supplementary Figure 1. All the DMN signals used in the current work were from the rats with electrodes in accurate DMN regions.

The dataset used in the current study was selected from the last 24 h of the total recording and was separated into three stages, including the resting (AWAKE), slow wave sleep (SWS) and rapid eye movement (REM) sleep stages. The rules for selecting each stage were based on LFP, EMG and videos, which have been summarized in our previous publications(Jing et al., 2017). Briefly, the scoring of the awake and sleep stages were performed by several experts. We included 29 rats in the current study. For each rat, 30 segments in different stages were chosen, and each segment lasted 10 s (a total of 300 s of LFPs).

SI Methods

**CAMP algorithm.** Using functional magnetic resonance imaging (fMRI) data, Liu and colleagues(Liu, Chang, & Duyn, 2013; Liu & Duyn, 2013) used the coactive pattern (CAP) method, a point process approach, to identify a set of CAPs with relevant network features to resting state networks, including the default mode network (DMN). In the present study, we developed a coactive micropattern (CAMP) measurement and specifically employed it to analyze neurophysiological data. An overview and procedure of the CAMP method is shown in Fig. 1. This method was used to extract CAMPs based on the extreme values of envelope signals at a high temporal resolution and reveal the fast dynamics of multichannel LFPs.

Several steps were involved in the CAMP method. First, the original data were bandpass filtered into specific frequency bands, such as the delta (1-4 Hz), theta (4-8 Hz), alpha (8-13 Hz) and beta (13-30 Hz) frequency bands. In our study, we filtered the original LFPs into the gamma (40-80 Hz) frequency band. Second, the Hilbert transform was applied to the filtered data to obtain the envelope signals (Fig. 2b). These envelope signals were normalized and downsampled (from 1,000 Hz to 100 Hz) to improve the signal-to-noise ratio (SNR) (Fig. 2c). The envelope signals were normalized by restricting the maximum value of the normalized envelope signal to 0.9 and the minimum value to 0.1. Third, the active points for each channel of envelope signals were then defined as the extreme points of the envelope signals, including local maximum and minimum values. Afterwards, the coactive patterns (CAPs) of the brain were introduced from normalized and down-sampled envelope signals for all states. The CAPs were the brain maps in which more than one brain region displayed active points at the same time point (Fig. 2d), and totally there were 1724646 CAPs that were extracted. Thus, we considered the number of brain regions with active points at the same time (parameter *N*) as one important parameter for extracting these coactive patterns.

After extracting the CAPs from envelope signals, we employed the k-means clustering algorithm to all the CAPs based on their spatial similarity to decompose the CAMPs (Fig. 2e). By clustering the CAPs into several distinct groups, we temporally divided the brain activity into multiple CAMPs. We repeated the k-means clustering with $k=2,\ldots, 10$ and applied the contour coefficient estimation (i.e., the sum of the squared errors, SSE) to determine the optimal number of distinct groups and select the optimal number of CAMPs. The optimal number of CAMPs was 3 in our study, according to the elbow of the curve between the *k* values and SSE values (Supplementary Figure 2b).

Next, the CAMPs were fit back to the coactive patterns (CAPs), assigning each CAP to the CAMP class with the lowest squared Euclidean distance to the three CAMPs. The CAMP index was then obtained from the assignment, which showed the temporal sequence of CAMPs in the gamma activity of the DMN. Then, we carefully updated the CAMPs and CAMP index according to the CAPs that belong to the same CAMP class using the following criterion:
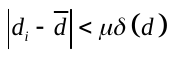
, where
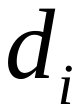
 is the squared Euclidean distance between the
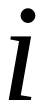
th coactive pattern with its assigned CAMP,
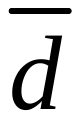
 is the mean of all distance,
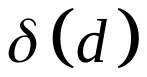
 is the standard deviation of these distance, and
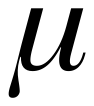
 is the penalty parameter we determined. If the distance did not conform to this criterion, then we removed the assignment of the corresponding CAPs from any CAMP class (Fig. 2f). Afterwards, we applied the k-means clustering algorithm to all remaining original CAPs using the same *k* value detected previously and redefined the clusters. This step was iterated until all distances obeyed this criterion. Using this criterion, we precisely determined the final spatial structures of CAMPs and the CAMP index. Notably, if the CAP was not assigned to any CAMP, we removed it from the CAMP index. 261344 CAPs were removed based on this criterion and there were 1463303 CAPs used in subsequent analysis. The final CAMP index only contained the assignments of all CAPs to their corresponding CAMPs.

In the present study, we calculated the number of CAPs for different N values ranging from 2 to 15 (Supplementary Figure 2a) and decomposed the CAMPs from these CAPs. Under different N values, the derived CAMPs exhibited similar spatial structures (Supplementary Figure 3). Therefore, considering the complexity of the calculation and requirement for additional information about DMN dynamics, we finally set N to 7 in the current study. Additionally, we also altered the penalty parameter
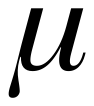
 from 1.5 to 3 (step size of 0.1) and described how the proportion of removed CAPs varied with different
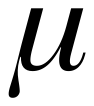
 values (Supplementary Figure 2c). By extracting the CAMPs with different
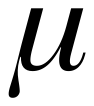
 values, we observed similar spatial structures of these CAMPs, implying that the penalty parameter might not affect the CAMPs (Supplementary Figure 4). Therefore, we finally set the value of this parameter to 2 in the subsequent analysis.

All our analyses of the CAMP algorithm were performed using our own custom MATLAB (release 2019a) scripts. If a special description was not included, we extracted the CAMPs of all the CAPs from whole segments acquired from rats in all three conscious stages. For every segment, the CAMPs were obtained by averaging the CAPs belonging to their corresponding clusters in that segment. Meanwhile, the CAMP index of each segment was also acquired from the total CAMP index.

**Estimation of the CAMP features.** In the present study, we employed five measurements to characterize the features of CAMPs and the CAMP index of each segment.

The total occurrence represented the number of CAPs assigned to each CAMP, and the occurrence probability was the proportion of the total occurrence of the number of all CAPs.

The total duration characterized the entire time required for each CAMP and the duration probability represented the proportion of that.

The duration of one CAP was defined as follows: the start time was the mid-point between the time point of this CAP and the preceding CAP, and the end time was the mid-point between the time point of this CAP and the next CAP.

We first defined the event for each CAMP to determine the mean duration of each CAMP. An event for each CAMP was that the coactive pattern before or after it should be different from itself. Thus, in the CAMP index, if the neighboring CAPs belonged to the same CAMP, then they should be included in one event for that CAMP. Using this approach, we obtained a new CAMP index in which the neighboring coactive patterns did not belong to the same CAMP. We separately estimated the numbers of events for all CAMPs, and the mean duration for each CAMP was calculated by dividing the total duration by the number of events for that CAMP.

All values of these CAMP features were calculated for each segment (10 s). The values of these features were averaged based on the rat and conscious stage to which they belonged to calculate the values of CAMP features for each rat in different conscious stages.

**Transition probabilities (TPs) for pairs of CAMPs.** The transition probabilities for pairs of CAMPs were the one-step and direct transitions among them. These TPs were separately estimated from the new CAMP index for each segment. Six types of direct transitions were identified in the new CAMP index. The TP for one direct transition was calculated by dividing the number of this transition by the total number of all direct transitions.

**Statistical analysis.** The statistical comparisons of the CAMP features and the TPs among CAMPs across the three stages were performed using the methods described below. First, an ANOVA was performed among all three stages, and then Student’s t test was performed as the post hoc test to determine the significance of differences between pairs of stages. In addition, the p values derived from Student’s t tests were corrected with the false discovery rate (FDR) correction.

**Reliability test for the three CAMPs across the 29 rats and different conscious stages.** We initially applied the CAMP analysis to the segments obtained from each rat in the AWAKE, SWS and REM sleep stages to assess the reliability of these CAMPs. Three different CAMPs were identified for each for each rat in each conscious stage (29*3*3 total CAMPs).

The reliability of CAMPs across rats was determined by estimating the correlation coefficients of the CAMPs among pairs of rats in the same conscious stage using the Pearson correlation method. These correlation coefficients were then averaged to obtain the reliability of each CAMP in each conscious stage (3*3). For different conscious stages, we next averaged the correlations across CAMPs and obtained the reliabilities of all CAMPs for each stage (Table 2).

For the analysis of the reliability of CAMPs across different conscious stages, we first estimated the correlation coefficients of CAMPs among pairs of conscious stages for each rat (29*3*3). Then, the whole correlation coefficient was averaged and the reliability of CAMPs across stages was obtained (3*3, Table 3).

**Randomization test for the CAMP index**. The randomization tests were applied to the new CAMP indices in the AWAKE, SWS and REM sleep stages(Lehmann et al., 2005). The null hypothesis was that if the transition from a preceding CAMP to the next CAMP occurred randomly, then the observed TPs would depend on the occurrence probability of CAMPs. During the test, we considered the expected TP from CAMP *X* to CAMP *Y* to be

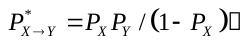
 (1)

where
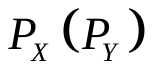
 is the occurrence probability for CAMP
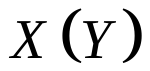
. The difference between the expected TP and the observed TP was then assessed by calculating the chi-square distance

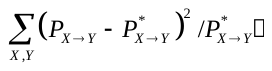
 (2)

where the sum was calculated for all 6 pairs of CAMPs for which
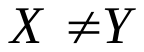
. The randomization test (permutation test) was then performed to statistically analyze the significance of this distance between the observed TP and expected TP. The permutation test was performed by shuffling the order of coactive patterns. The number of randomizations in the permutation test was set to 10,000 in our study, and the probability was determined by the rank of the observed difference among the randomly obtained differences.

**Phasic relationships between CAMPs and up-down states in the slow oscillations of the DMN during deep sleep.** We first averaged the DMN activity to obtain the activity of anterior DMN and posterior DMN in the SWS stage and to assess the phasic relationship between CAMPs and up-down states. The average activity was then bandpass-filtered at 0.5-2 Hz and downsampled from 1,000 Hz to 100 Hz, which coincided with the CAMP algorithm. Using this approach, we finally obtained the downsampled slow oscillations in the anterior DMN and posterior DMN regions. Then, the Hilbert transform was applied to these slow oscillations to obtain the instantaneous phase for both slow activity in the anterior DMN and posterior DMN activity. By combining the acquired instantaneous phase and timing of each CAMP, we obtained the instantaneous phases of all CAMPs in the slow activity of the anterior DMN and posterior DMN, and the distributions of phases for the cDMN, aDMN and pDMN in the SWS stage. Finally, we employed the Rayleigh test to analyze the non-uniformity of these distributions of phases for the three CAMPs.

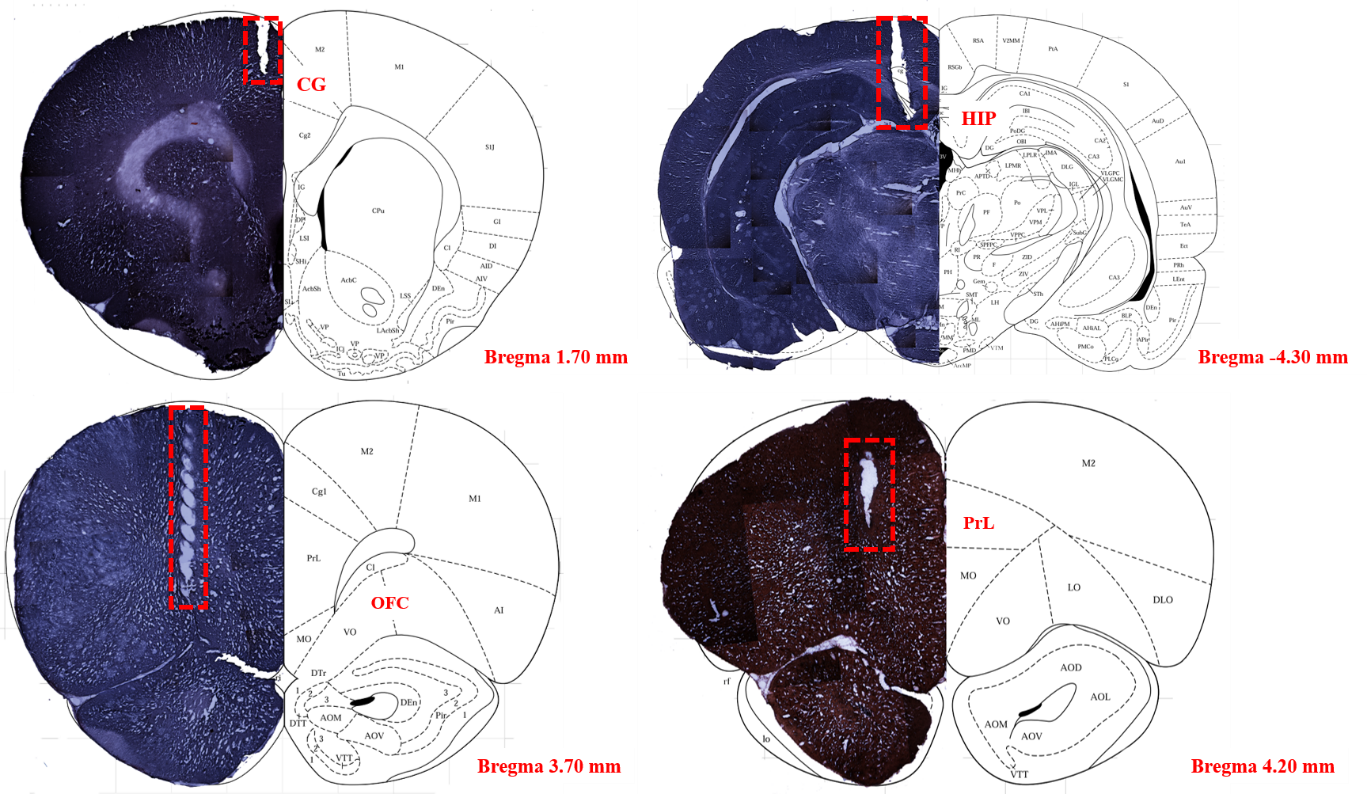

**Supplementary Fig. 1.** The histological images for four depth regions (eight depth electrodes), including the PrL, OFC, CG and HIP regions.

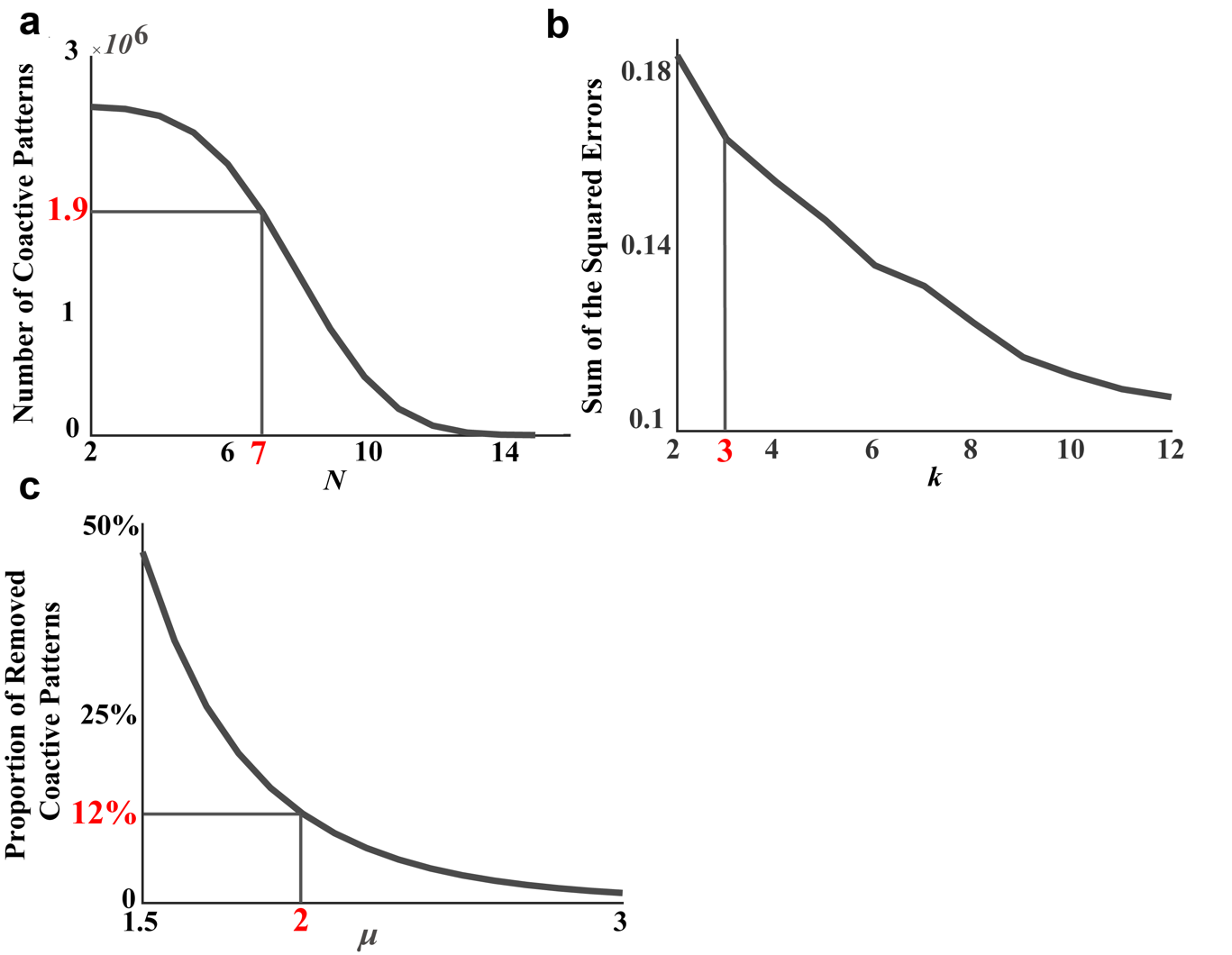

**Supplementary Fig. 2**. The methods used to select the number of coactive brain regions, optimal numbers for groups and
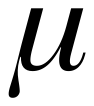
. (a) The relationship between the number of coactive points and the number of coactive brain regions (*N* from 2 to 15, step size of 1). (b) The relationship between k clusters (from 2 to 12, step size of 1) and SSE to choose the best k clusters. (c) The relationship between
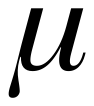
 (from 1.5 to 3, step size of 0.1) and the proportion of removed CAPs.

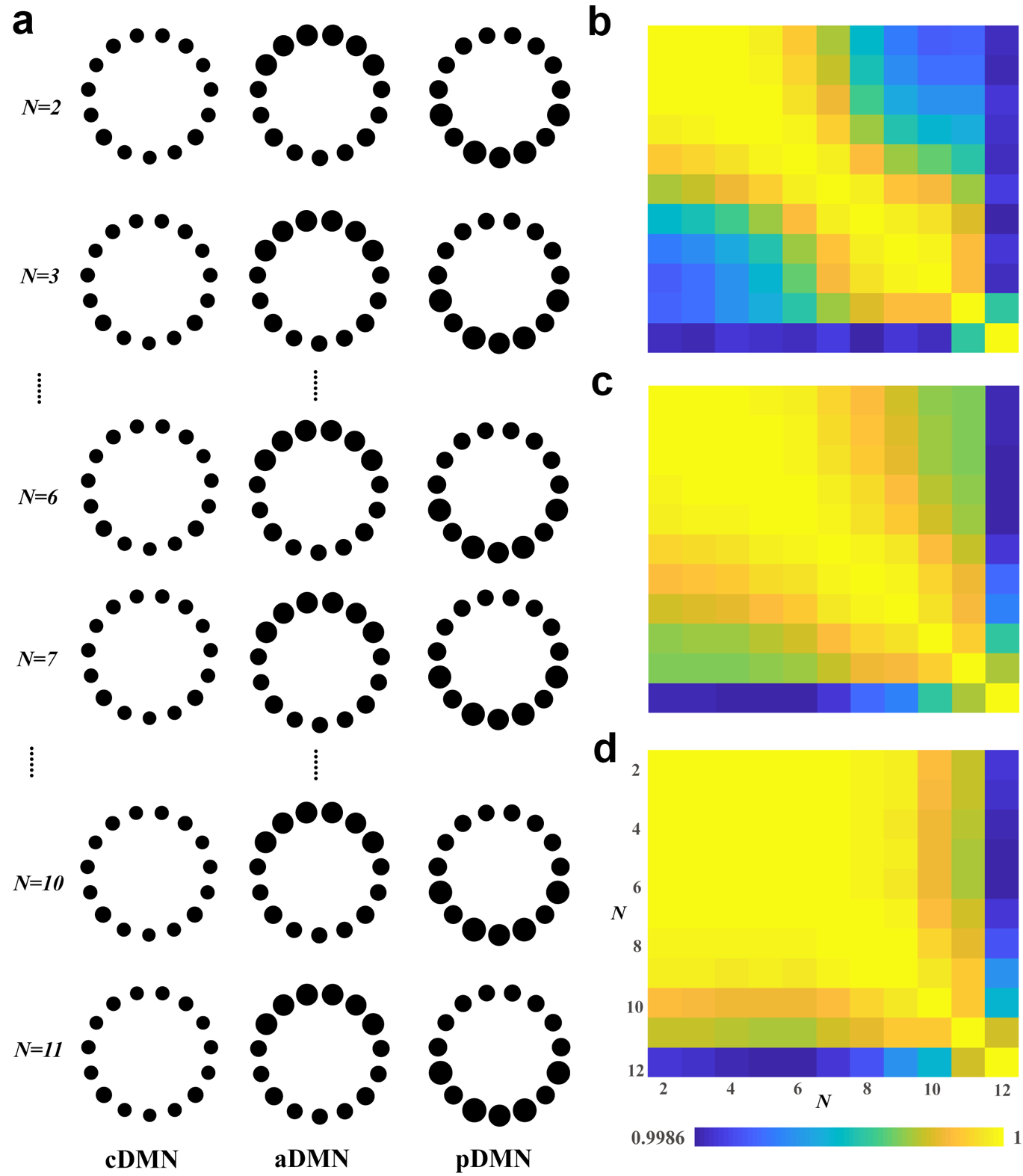
 **Supplementary Fig. 3**. The effect of N (number of coactive brain regions) on the CAMPs. (a) The CAMPs in different N (N ranges from 2 to 11). (b) The structural correlations among the cDMN in different N. (c) The structural correlations among the aDMN in different N. (d) The structural correlations among the pDMN in different N.

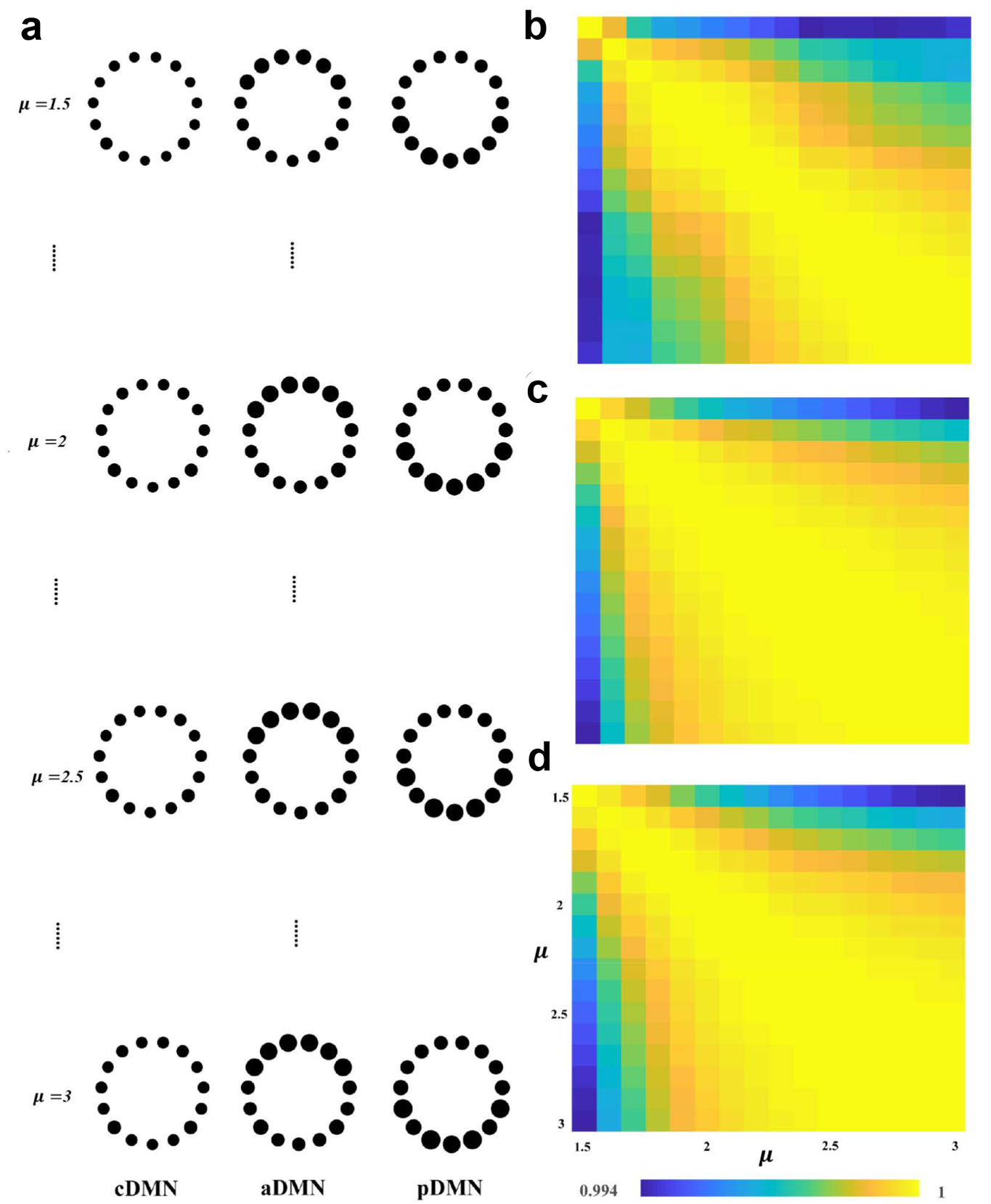

**Supplementary Fig. 4**. The effect of
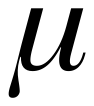
 on the CAMPs. (a) The CAMPs at different
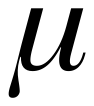
 values (
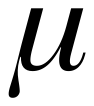
 ranges from 1.5 to 3). (b) The structural correlations among the cDMN with different
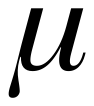
 values. (c) The structural correlations among the aDMN with different
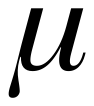
 values. (d) The structural correlations among the pDMN with different
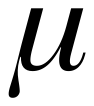
 values.

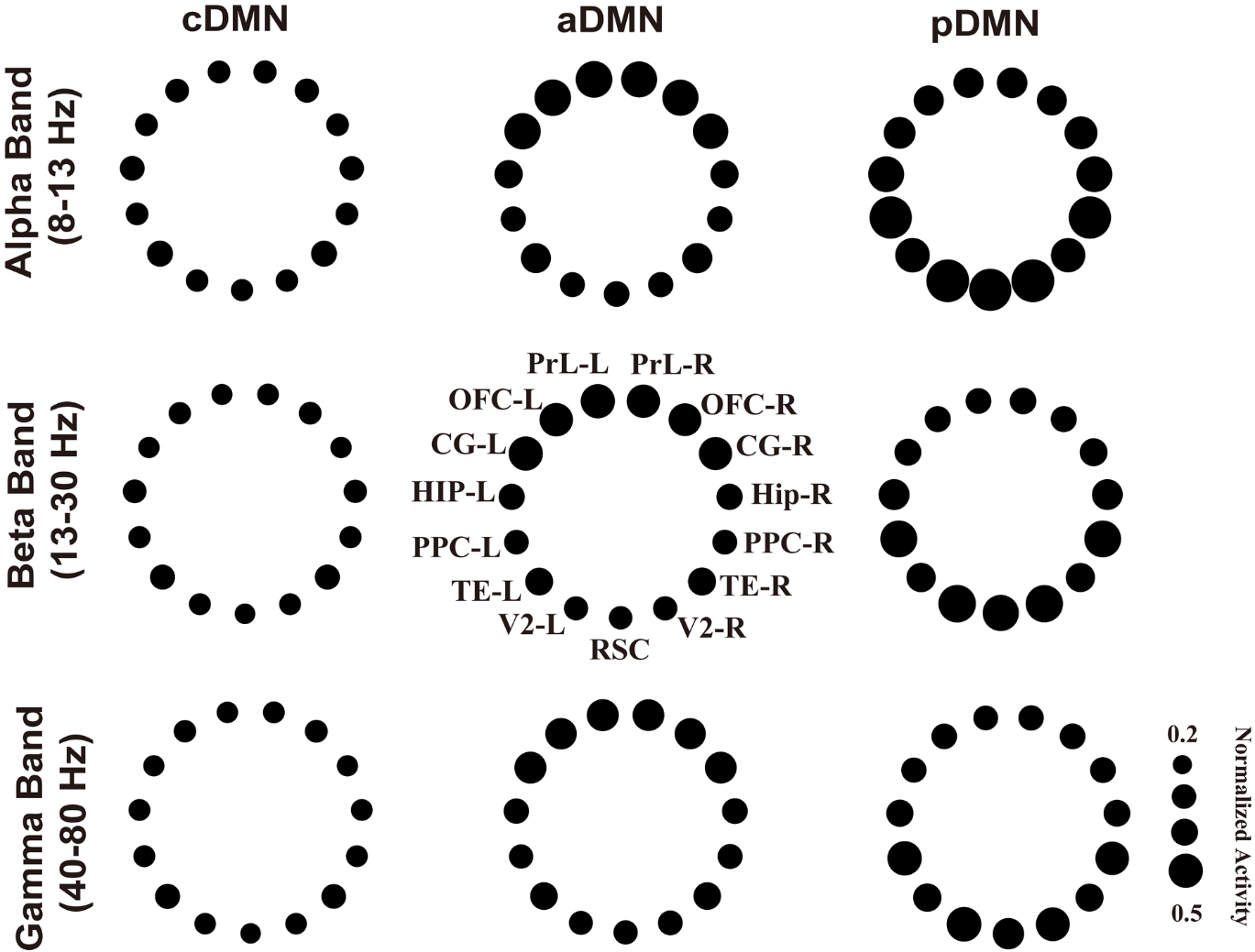

**Supplementary Fig. 5.** The CAMPs derived from DMN alpha activity (8-13 Hz), beta activity (13-30 Hz) and gamma activity (40-80 Hz) during wakefulness-sleep cycle. We observed similar spatial structures for cDMN, aDMN and pDMN among three kinds of DMN activity.
